# POLD replicates both strands of small kilobase-long replication bubbles initiated at a majority of human replication origins

**DOI:** 10.1101/174730

**Authors:** Artem V. Artemov, Maria A. Andrianova, Georgii A. Bazykin, Vladimir B. Seplyarskiy

## Abstract

Error-prone mutants of polymerase epsilon (POLE*) or polymerase delta (POLD1*) induce a mutator phenotype in human cancers. Here we show that the rate of mutations introduced by POLD1* is elevated by 50%, while the rate of POLE*-induced mutations is decreased twofold, within one kilobase from replication origins. These results support a model in which POLD1 replicates both the leading and the lagging strands within a kilobase from an origin. The magnitude of the mutational bias suggests that the probability of an individual origin to initiate replication exceeds 50%, which is much higher than previous estimates. Using additional data from nascent DNA sequencing and Okazaki fragments sequencing (OK-seq) experiments, we showed that a majority of origins are firing at each replication round, but the initiated replication fork does not propagate further than 1Kb in both directions. Analyses based on mutational data and on OK-seq data concordantly suggest that only approximately a quarter of fired origins result in a processive replication fork. Taken together, our results provide a new model of replication initiation.

---

Accumulation of sequencing data is improving the understanding of processes involved in mutagenesis. Recently, analysis of preferential fork direction helped uncover a major mode of APOBEC-induced mutagenesis in cancer (Seplyarskiy et al. 2016a; Morganella et al. 2016; Haradhvala et al. 2016) and revealed the dominant role of mismatches introduced by POLD1 in mutagenesis of cancers with mismatch repair deficiency (Andrianova et al. 2017). Understanding of how major DNA polymerases, POLE and POLD1, divide the labor during DNA replication has largely stemmed from patterns of mutation or ribonucleotide incorporation in cell systems with deficiencies in different proofreading mechanisms (Nick McElhinny et al. 2008; Larrea et al. 2010a; Lujan et al. 2012; Johnson et al. 2015a; Reijns et al. 2015; Clausen et al. 2015; Andrianova et al. 2017). By contrast, a qualitative model of replication has been based on reconstruction of replication *in vitro* (Yeeles et al. 2015; Georgescu et al. 2014; Yeeles et al. 2017; Kurat et al. 2017; Devbhandari et al. 2017), and efficiency of replication origins in mammalian cells has been estimated mainly from sequencing of 500-2500 nucleotide long stretches of nascent DNA (Cayrou et al. 2011; Besnard et al. 2012, 2014; Cayrou et al. 2015). Analyses of mutagenesis in systems with deficiencies of different components of proofreading machinery have not yet been applied to study firing of replication origins. Here, we use somatic mutations in cancers as well as OK-seq and repli-seq data to qualitatively and quantitatively revisit the model of origin firing.

Mutational traces of low-fidelity POLE (POLE*) or POLD1 (POLD1*) mutants may be used to trace the genomic regions replicated by each polymerase. However, mismatches introduced by polymerases are co-replicatively repaired by mismatch repair system (MMR), which may bias the mutational traces left by POLE* or POLD1*. This bias would not exist in tumours with biallelic mismatch-repair deficiency (bMMRD) or MMR deficiency manifested as microsattelite instability (MSI). In our analyses we used somatic mutations collected by whole-genome sequencing of POLE* MSI endometrial tumours and by whole-exome sequencing of POLD* bMMRD glioblastoma samples.

ORC binding sites determined by ChIP-seq in HeLa cells (Dellino et al. 2013a) were taken as markers of potential origins. To obtain a mutation-based model of replication, we estimated context-corrected mutation rate for the 5 kb around each ORC site (see Methods). We discovered a twofold drop of mutation rate within a kilobase from ORC sites in POLE* MMR-deficient tumours (*P* = 1.2∗ 10^−157^, Figure 1A). The drop had a similar magnitude for different substitution types (Figure S1A) and for genomic contexts excluding CpG dinucleotides (Figure S1B). In contrast to POLE* MMR-deficient cancers, POLD* MMR-deficient tumours possessed a 1.5-fold higher mutation rate nearby ORC sites as compared to the flanks 5Kb apart (*P* = 2.3 ∗ 10^−4^), (Figure 1A). To control that a mutation rate peak at replication origins in POLD* tumours could not be explained by biases in genomic distribution specific to exome data, we studied POLE* tumours for which only whole-exome data were available. No peak was observed around replication origins (Figure S2). Replication origins are characterized by a very specific epigenetic profile, and to account for this, we plotted average mutation rates around DNA features which have been shown to affect the local mutation rate: gene transcription start sites (Sabarinathan et al. 2016; Perera et al. 2016), the peaks of H3K4me3, H3K9ac, H3K27me3 histone marks, binding sites of CTCF, cohesin (SMC3) (Schuster-Böckler, Benjamin, and Ben 2012); and SUZ12 binding sites that were believed to be associated with replication process (Cayrou et al. 2015). ORC sites and SUZ12 had the strongest effect on local mutation rate (Figure 1B). We performed multiple regression predicting mutation rate in 1kb genomic windows based on epigenetic factors and showed that the presence of an ORC site was the best predictor of the local mutation rate in POLE* tumours (*P*_*ANOVA*_ < 2∗10^−16^). Epigenetic background, such as DNAse sites which frequently overlapped with ORC sites, could not explain the effect of ORC sites on mutation rates in POLE* and POLD1* tumours (Figure S3A,B). We also controlled for replication timing and preferential fork direction (Figure S4A), but did not find any decrease or increase in mutation rate for a control set of genomic regions (Figure S4B). Therefore, the effect was associated with origins themselves and could not be explained by clustering of origins in the domains of early replication timing. We also explored APOBEC-induced mutational patterns, which had been recently linked to replication (Haradhvala et al. 2016; Seplyarskiy et al. 2016b), but did not see any alterations of mutation rate near ORC sites (Figure S5). The observed accumulation of mutational trace left by POLD and the depletion of POLE-introduced mutations at ORC sites suggests that POLD replicates both the leading and the lagging strands 1kb from the origin site, while the two polymerases canonically divide the labor further apart from the origin: POLE replicates the leading strand whereas POLD replicates the lagging strand (Figure 1C).

POLE* and POLD1* were known to introduce mutations in different trinucleotide contexts (Andrianova et al. 2017; Shlien et al. 2015). If context preferences may be extrapolated on non-mutated polymerases in systems with dominant contribution of polymerase errors to mutagenesis, we may use somatic mutations in specific contexts to trace the contribution of each polymerase. Mutation rate in many nucleotide contexts in MMR-deficient cancers is about 10 fold higher than that in tumours with functional MMR (Andrianova et al. 2017) and this increase can be primarily attributed to unrepaired mismatches introduced by replicative polymerases. In line with our model, in MMR-deficient cancers with non-mutated replicative polymerases, mutation rate was twofold decreased in POLE*-associated contexts (Gp<u>C</u>pG→T and Cp<u>G</u>pC →A) at ORC sites (Figure S6, *P* = 2∗ 10^−16^), whereas the difference was very minor for POLD1*-associated context (Gp<u>C</u>pC→T and Gp<u>G</u>pC→A, *P* = 0.15).

**Figure 1.**
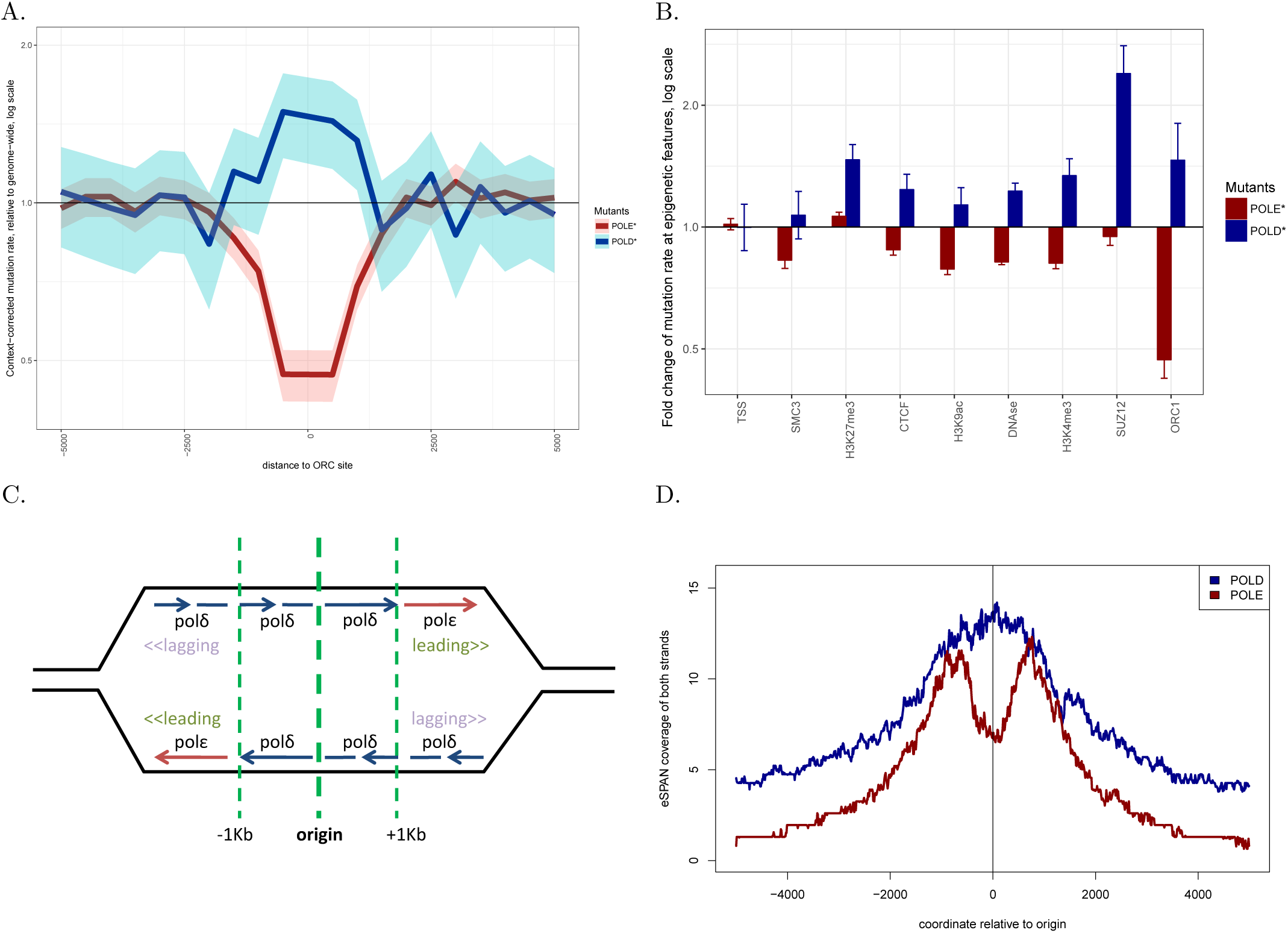
Local somatic mutation rates on a 5-kilobase scale centered around various epigenetic features in POLE* tumours and in POLD* tumours. Note that only exome data were available for polD deficient tumours. (A) For each epigenetic feature (x-axis), we plotted context-corrected mutation rate adjacent to this feature (-500..+500bp window) normalized by mutation rate further from the regions of interest (-5000..-4500 bp and 4500..5000 regions, see Methods section). ORC1 sites were associated with the strongest drop in local mutation rate in POLE* tumours and, in the same time, with a strong and significant increase of mutation rate in poD-deficient tumours. (B) Context-corrected somatic mutation rates (see Methods) on a 5-kilobase scale centered around potential human replication origins defined as ORC binding sites in POLE* tumours and in POLD* tumours. Note that only exome data were available for POLD* tumours.

Additionally to human data, we studied the genomic distribution of POLE and POLD1 binding with respect to positions of replication origins in *Saccharomyces cerevisiae* (Yu et al. 2014). We reanalyzed the available ChIP-seq based eSPAN data and plotted median profiles of PolE and PolD binding around replication origins (Figure 1D). In fact, while binding of both PolD and PolE was increased within ∼ 5Kb of yeast replication origins, PolE binding was significantly decreased at the ∼1Kb around them, whereas PolD binding was increased here. This result in *S. cerevisiae* was in agreement with the result of a different polymerase tracing method based on ribonucleotide incorporation in *Schizosaccharomyces pombe* (Daigaku et al. 2015). Therefore, PolE depletion at origins in yeast was confirmed by a method that was independent of mutagenesis.

Similarly to the results obtained for human tumours, we observed a 1.5-fold increase of mutation rate at origins in *pol3-L612M* (POLD*) *Saccharomyces cerevisiae* strain (Lujan et al. 2014), (Figure S7). We did not observe a significant drop of mutation rate in *pol2-M644G* (POLE*) yeast strain likely due to high noise rate. These results show that the replication programme involving DNA synthesis near origins by POLD both on the leading and the lagging strands is conserved between humans and yeast. Our yeast results are in line with experiments in yeast reconstructing a processive replication fork *in vitro* which required synthesis of both strands by POLD in the early stage of replication (Yeeles et al. 2017; Kurat et al. 2017; Devbhandari et al. 2017).

Such strong and reverse local mutation rate changes near ORC sites in POLE* MMR-deficient and POLD1* MMR-deficient cancers not only suggested a specific mode of replication at origins, but also implied very efficient firing of origins. Indeed, even if we conservatively assumed that replication of both strands near a fired origin did not involve POLE at all, a twofold decrease in mutation trace of POLE would mean that at least half of all origins initiated replication (see Methods). Moreover, a 1.5-fold increase of the mutation rate in POLD* MMR-deficient cancers again means that half of origins fire (see Methods). This conservative mutation-based estimation was in a sharp contrast to the results obtained by other methods, which estimated the probability of an individual origin firing in mammalian cells to reside in the range of 10-30% (Löb et al. 2016; Fragkos et al. 2015; Cayrou et al. 2015). We inferred similar estimations of origin firing probability between 0 and 30% (95% confidence interval) from asymmetry of POLE*-induced mutations within 5kb from ORC sites (Figure S8A). Asymmetry of APOBEC-induced mutations also suggested origin efficiency of 0-20% (95% confidence interval, Figure S8AB). To resolve this contradiction, we hypothesized that most potential origin sites marked by ORC binding initiate replication, while only a fraction of the forks proceed further than 1Kb (Figure 2).

We tested this hypothesis with additional data on nascent DNA sequencing from repliSeq experiments (Hansen et al. 2010) and from Okazaki fragment sequencing (OK-seq), (Petryk et al. 2016).

Our analysis of human repliSeq data proved the presence of 1Kb-long replication forks at many ORC sites. We stratified all ORC sites into four groups according to their replication timing. Early origins in the late G1 phase and the early S phase showed a twofold increase in the normalized content of *de novo* synthesized DNA (Figure 3A).

**Figure 2.**
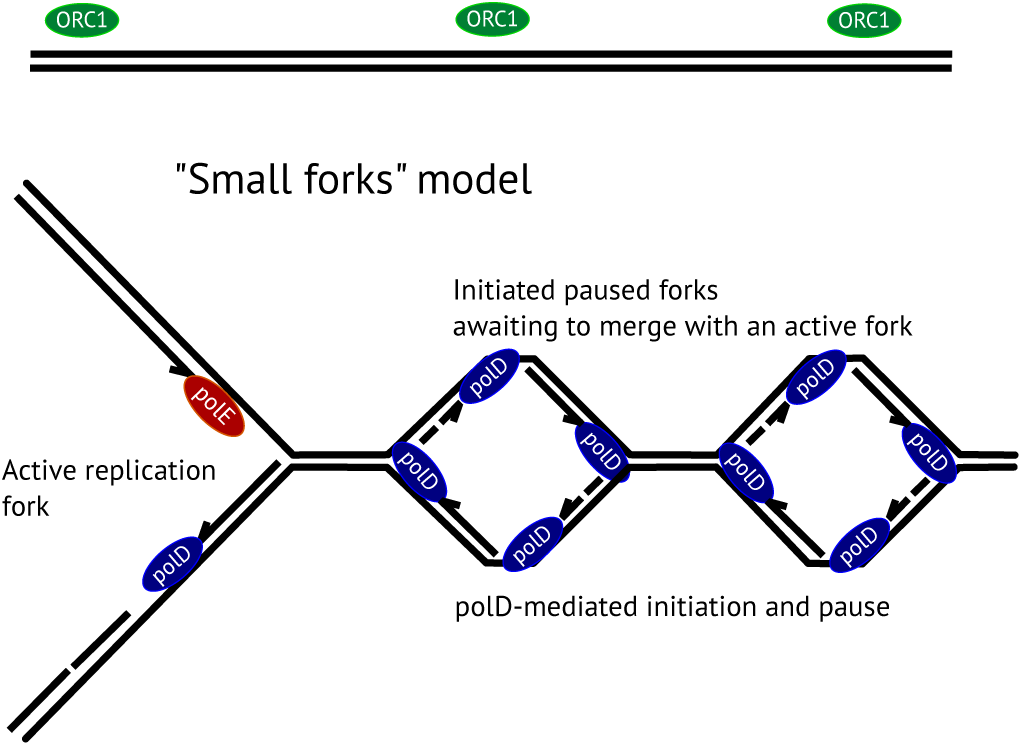
Model: replication is initiated by POLD even at dormant origins, however, most of the produced forks pause soon after initiation and await to be merged with an active replication fork.

Moreover, a 1kb-wide peak around ORC sites suggested that the forks at most ORC sites indeed did not propagate further than 1kb: otherwise, we would expect a processive fork to move 100Kb from an initiation site (Hansen et al. 2010) making it impossible to observe any local effect on a kilobase scale.

Being aimed to sequence Okazaki fragments, OK-seq captures *de novo* synthesized DNA fragments shorter than 200bp. Therefore, both Okazaki fragments and short stretches of nascent DNA contribute to OK-seq signal. Further than a kilobase from an origin, OK-seq captures only Okazaki fragments that reflect strand-specific replication. Therefore, firing probability of an origin can be estimated from the difference between OK-seq signal to the left and to the right of the origin on the Watson and on the Crick strands (see Methods). The 11% average difference between OK-seq signal on the Watson and the Crick strand 5kb from ORC sites reflects the preferential fork direction and yields the probability of an individual origin to initiate a processive replication fork of 11%. On top of that, we observed a 1.5-fold enrichment of OK-seq reads within 1kb from ORC sites compared to mean OK-seq signal 5kb from ORC sites, which is 4.4-fold stronger than the level of asymmetry 5kb apart from an ORC sites (Figure 3B, Figure S9). This likely indicates the presence of paused replication forks. Therefore, for an ORC site there is a 11% chance to initiate a processive replication fork and a 48% chance to initiate a fork which is paused 1kb from the initiation site. This gives a total chance of origin firing of 59%, which is very close to mutation-based estimations.

Previous studies could underestimate the chances of an individual origin to initiate replication as they were aimed to look for processive replication forks and operated on 10kb - 1Mb genomic scale. In this work, we studied human replication on a kilobase scale with several independent methods and showed that origin firing probability was higher than it had been previously expected, though most of the initiated forks were paused at a distance less than 1kb from the initiation and did not proceed further awaiting to be merged with a processive fork. Such small replication forks are barely detectable by nascent DNA sequencing used for origin mapping, because considered DNA fragments were longer than 1 KB (Besnard et al. 2012). However, nascent DNA fragments shorter than 200 nucleotides were detectable in OK-seq and in repliSeq data which allowed us to observe small paused replication forks. Our results could also settle a debate between (Johnson et al. 2015b) and (Lujan et al. 2014; Larrea et al. 2010b;

Burgers, Gordenin, and Kunkel 2016; Watt et al. 2016; Lujan, Williams, and Kunkel 2016) on the roles of POLE and POLD in replication of the leading and the lagging strand. In (Johnson et al. 2015b), a study in a yeast model suggested that POLD should replicate both the leading and the lagging strand. Here we show that this claim would be true within a kilobase around a replication origin, which was exactly the size scale of the region where mutations were counted in the original study. In contrast, on a distance of 10-100 Kb, an asymmetric mutational pattern (Figure S10) can be explained by differential replication of the leading and the lagging strand by POLE and POLD respectively (Lujan et al. 2014; Larrea et al. 2010b). We hope that the new model of replication will be helpful for the analyses of somatic mutagenesis in cancer and will stimulate additional experimental efforts to understand molecular machinery which orchestrates this behavior of replication origins.

**Figure 3.**
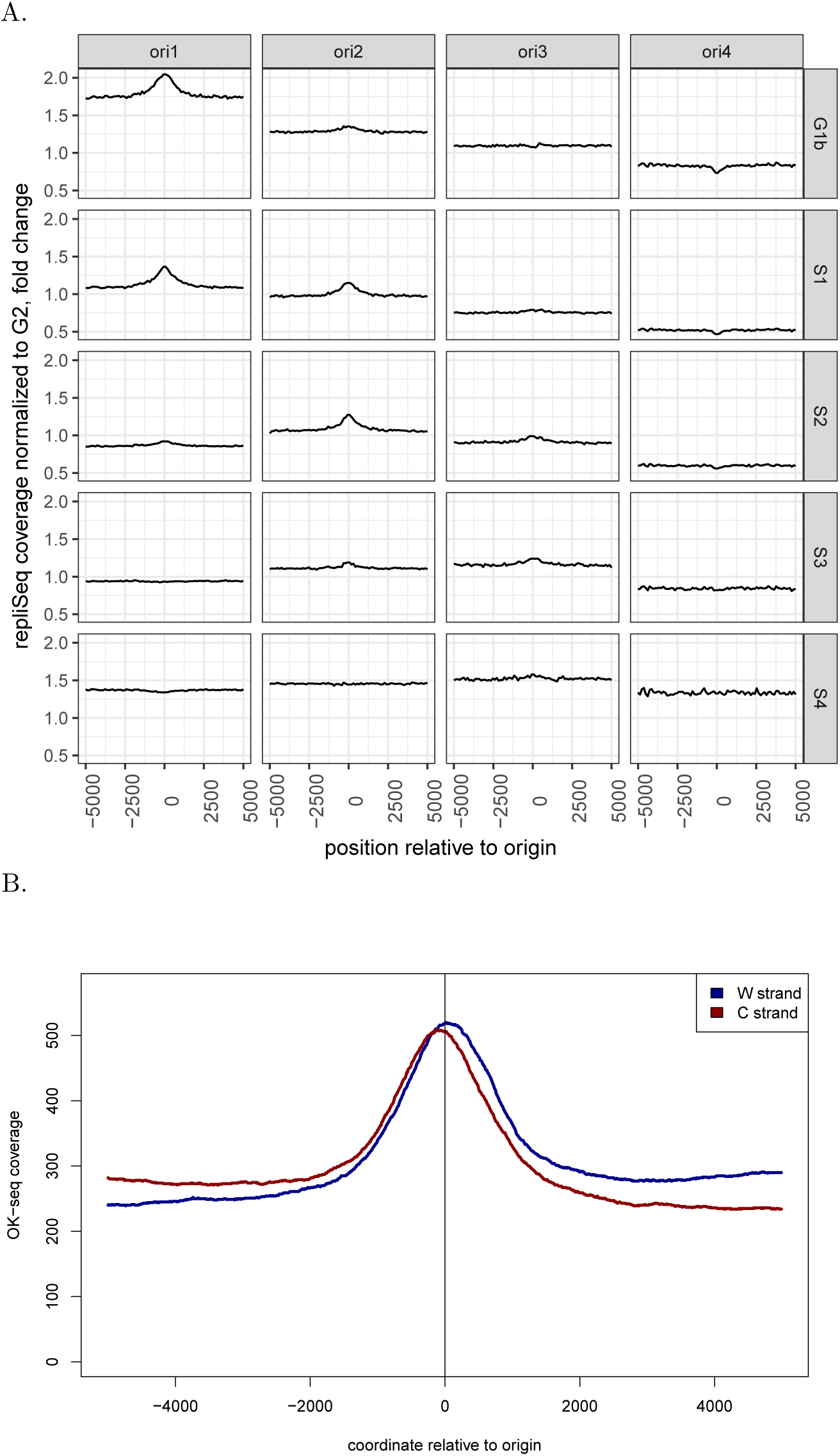
(A) Mean repliSeq profile at ORC sites. repliSeq signal at G1b, S1, S2, S3 and S4 phases of the cell cycle was normalized to the total number of reads in a sample and to the signal at G2 phase where all DNA was expected to have equal copy number. ORC sites were split into four groups according to their replication timing. (B) Density of Okazaki fragments profiled by OK-seq around ORC sites. The fragments mapped to Watson (W) and Crick (C) strands are plotted separately. The enrichments of short *de novo* synthesized DNA fragments at ORC sites exceeds the difference of Okazaki fragment density between the leading and the lagging replication strands.

## Online methods

Locations of human replication origins were taken from an experiment in which Orc1 ChIP-seq was combined to replication timing profiling (GEO accession number GSE37583), (Dellino et al. 2013).

Localization of ChIP peaks for histone marks and SUZ12 was obtained from ENCODE project. TSS were fetched from ENSEMBL genome annotation.

As DNAse hypersensitive sites showed decreased mutability in POLE* cancers, we controlled for DNAse accessibility within ORC1 binding sites. Indeed, 4697 (or 82%) ORC sites overlapped with a DNAse hypersensitivity site, moreover, those DNAse sites showed greater occupancy score than did an average DNAse site. Therefore, we constructed a control set of DNAse sites which had the score distribution similar to that of the DNAse sites overlapping with ORC1 binding sites (Figure S2). For the control set of DNAse sites we estimated mutability in POLE* and POLD* cancers, but the effect magnitude didn’t exceed that of ORC binding sites.

The data on whole-genome cancer somatic mutations were obtained from TCGA. The annotation of TCGA tumours with defunct polE or defunct MMR (MSI) was taken from (Shinbrot et al. 2014). Whole-exome data for tumours with mutant polE and polD was taken from (Shlien et al. 2015).

Distribution of mutation rates for a set of genomic loci of interest (e.g., replication origins) was plotted as follows. For each locus of interest, it’s surrounding regions (e.g., 5Kb upstream and 5Kb downstream) were each split into 10 consecutive bins of equal length (in other words, 20 bins were analyzed for each locus). In each bin, we calculated the number of mutation in each context or the total number of mutation. By context we mean a particular substitution (1-nucleotide context, e.g. C→T). The only exception was the case of APOBEC-induced mutations for which 3-letter contexts were considered. APOBEC mutations on forward strand were defined as TCW→ K, APOBEC mutations on reverse strand were defined as WGA→M (where W = A or T; K = G or T; M = A or C). We also calculated nucleotide content in each bin. In the case of APOBEC, the number of potentially mutagenic contexts (TCW and WGA) was calculated. Next, we aggregated the obtained measures for each relative bin position over all loci of interest. Mutation rates were calculated either for all substitutions (as in Figure 1) or for each possible substitution separately. Error bars on the plot depict standard deviation for the calculated mutation rate assuming that number of mutation was Poisson-distributed.

To calculate context-corrected mutation rate in each bin, we first fetched trinucleotide contexts for all studied mutations in a certain group of tumours. For each trinucleotide context (e.g., AAG), we calculated its mutability as the number of mutations observed in this context (only mutations in the second nucleotide were considered) normalized to the total number of instances of the context in the genome. For each bin, we calculated the number of each trinucleotide contexts and aggregated those numbers according to bin position relative to a region of interest (e.g., a total number of contexts AAG in bins -500..0 from all ORC sites). Expected number of mutations was calculated as a sum of trinucleotide counts multiplied by estimated trinucleotide mutability over all trinucleotide contexts:

Σ _*XYZ*_ *Mutability*(*XYZ*) ∗ *Count*(*XYZ*), where X, Y and Z ⊂ {*A, C, G, T*}. Finally, the observed number of mutations for each relative bin position was normalized to its expected value. For plots 1A and 1B, we normalized context-corrected mutation rate in each bin to the mean rate obtained for the furthermost bins (i.e., bins -5000..-4500 and +4500..+5000).

Mutation rates were calculated as a count of mutations in a particular context in a genomic bin divided by the number of times this nucleotide context appeared in a given bin.

Magnitude of the drop in mutation rate in POLE* tumours was used to calculate the probability of origin firing as follows. Let *p* be the average probability of firing of an individual origin and *r*_*E*_ be the average mutation rate of POLE per strand. Far from origins, POLE is believed to replicate the leading strand whereas POLD replicates the lagging strand, therefore, we expect a genomic average rate of POLE mutagenesis to be *r*_*E*_. To provide the lower estimate for p, we assume that both strands near an active origin are replicated only by POLD. In *p* cases, the corresponding locus would acquire 0 mutations introduced by POLE whereas in (1 − *p*) cases the origin would not fire and the locus would acquire a genomic-average number of mutations (*r*_*E*_). Therefore, *p* is exactly the size of the drop in POLE mutagenesis. Thus, for *p* = 0.5, a 50% drop in mutation rate in POLE* tumours is expected.

For the mutations introduced by POLD, *p* can be derived in the following way. Let *r*_*D*_ be the average mutation rate of POLD per strand. As POLD is believed to replicate only the lagging strand far from origins, *r*_*D*_ would be an average rate of POLD mutability. In *p* cases (origin fired), an origin locus is replicated exclusively by POLD and acquires mutations with 2 ∗ *r*_*D*_ rate, whereas in (1 − *p*) cases (origin did not fire) the rate would be *r*_*D*_ as only one strand is replicated by POLD. This yields the expected mutation rate at origins of 2∗ *p* ∗ *r*_*D*_ + (1*−p*)∗*r*_*D*_ = (*p* + 1) ∗ *r*_*D*_. Therefore, a relative drop of POLD mutagenesis at origins as compared to genomic average would be (*p* + 1) *r*_*D*_*/r*_*D*_ = (*p* + 1). Thus, for *p* = 0.5, the mutation rate in POLD* tumours is expected to be (0.5 + 1) = 1.5 times higher at ORC sites than the genomic average.

To estimate origin firing probability from mutational asymmetry, we assumed that if the replication fork passed a certain genomic locus always in one direction (e.g., to the right), it would leave *α* times more mutations on the leading strand (e.g., *C*→*A*) than on the lagging strand (e.g., *G*→*T*). We took a conservative estimate of *α* = 3 for *C*→*A/G*→*T* pair of substitution (Shinbrot et al. 2014). Let *f* be the probability of a fork to traverse a certain locus from the left to the right in an act of replication (1 − *f* being a probability of the fork to move in the opposite direction), and let *K* be the ratio between the rates of complementary mutations in a locus (e.g., 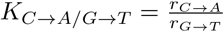).

For an arbitrary *f*, 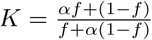. Note that *f* = 1 yields *K* = *α* as expected by the definition of *α*. An upper estimate for *K* was calculated from experimental data on mutational asymmetry as the ratio between complementary mutations plus 1.95 ∗ *sd* of that ratio giving the upper bound of a 95% confidence interval (e.g.,*KC*→A/G→T ≈ 1.3). Simple derivations give the expression for 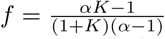. Let *p* be the probability of an origin to fire. If we consider a region to the right of an origin, e.g. (+1kb..+5kb), in *p* cases the fork will proceed this region from the left to the right and in 1 − *p* cases it will proceed in a random direction. Thus, the probability *f* of a fork to traverse the region from the left to the right 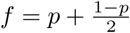, which is equivalent to *p* = 2*f* − 1. Finally, we can estimate the average origin firing probability *p* from mutational asymmetry 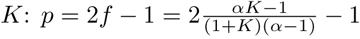. For a particular case of *α* = 3 and *K*_*C*→*A/G*→*T*_ ≈1.3 this yields *p* ≈ 0.26.

The following procedure was applied to estimate the probabilities of initiating a processive replication fork and a paused replication fork at an origin from OK-seq data, Let *c*_*-∞*_, *c*_0_ and *c*_*∞*_ be the averaged values of OK-seq signal for the Crick strand 5kb to the left from origin, at origin and 5kb to the right of origin, respectively. Let *w*_*-∞*_, *w*_0_ and *w*_*∞*_ be similar values for the Watson strand. Genomic-averaged OK-seq signal for the Watson and the Crick strands were estimated as 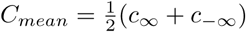 and 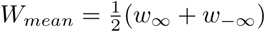 respectively. The relative difference between OK-seq signal on each strand to the left and to the right of an origin (i.e., 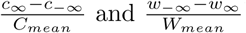) both estimate the probability *p*_*processive*_ of an origin to initiate a replication fork which propagates further than 1kb. To estimate the fraction of replication forks which initiated and did not proceed further than 1Kb, we calculated the excess OK-seq signal at origin relative to genomic average OK-seq signal (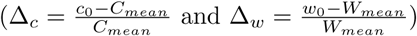). Δ_*c*_ and Δ_*w*_ were compared to 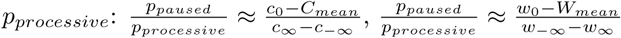.

## Supporting Information

**Figure S1.**
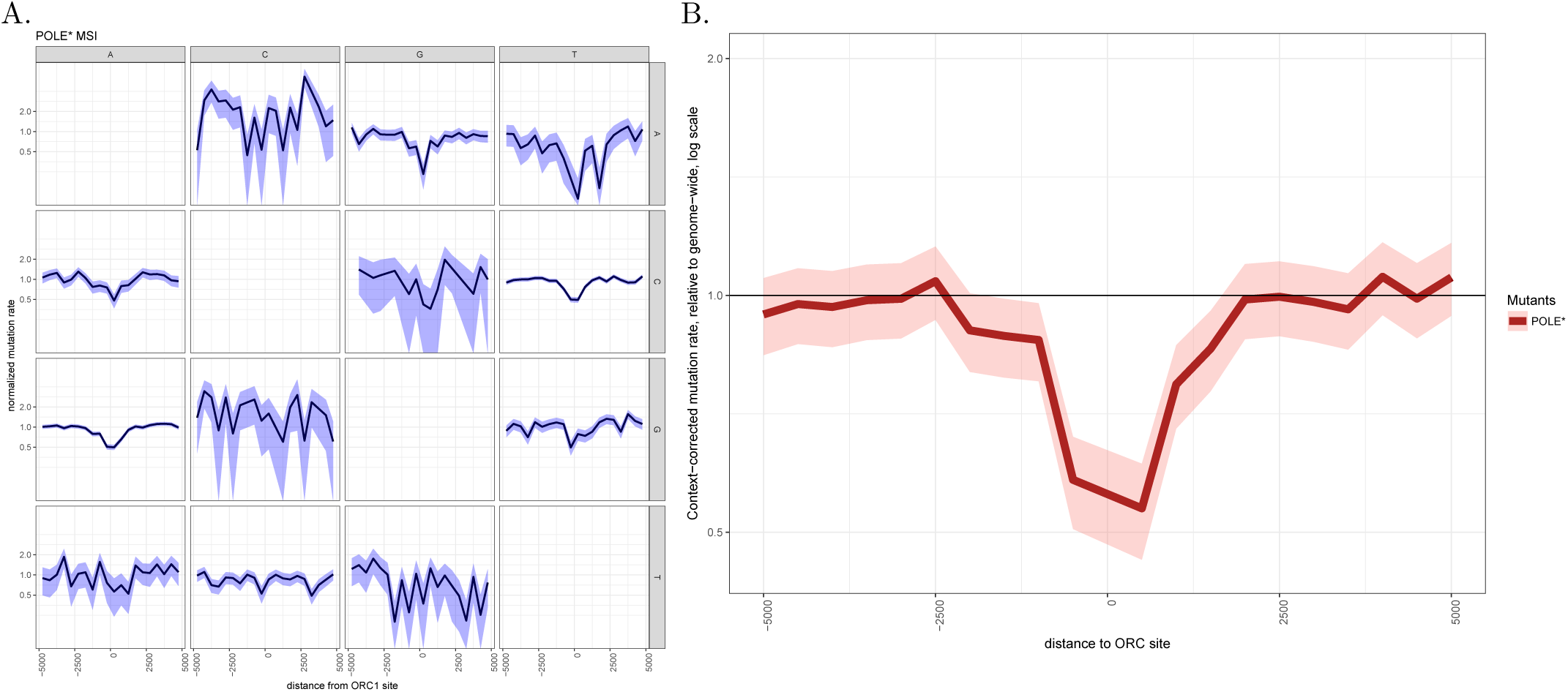
(A) Somatic mutation rates on a 5-kilobase scale centered around potential human replication origins defined as ORC binding sites in POLE* tumours calculated independently for each possible substitution and normalized to nucleotide content (see Methods). A letter which mark a row represents the reference nucleotide, a letter which marks a column represents alternative somatic variant observed in the tumour. (B) Similar to Figure 1A for non-CpG contexts. Context-corrected somatic mutation rates (see Methods) on a 5-kilobase scale centered around potential human replication origins defined as ORC binding sites in POLE* tumours for nucleotide contexts excluding CpG contexts.

**Figure S2.**
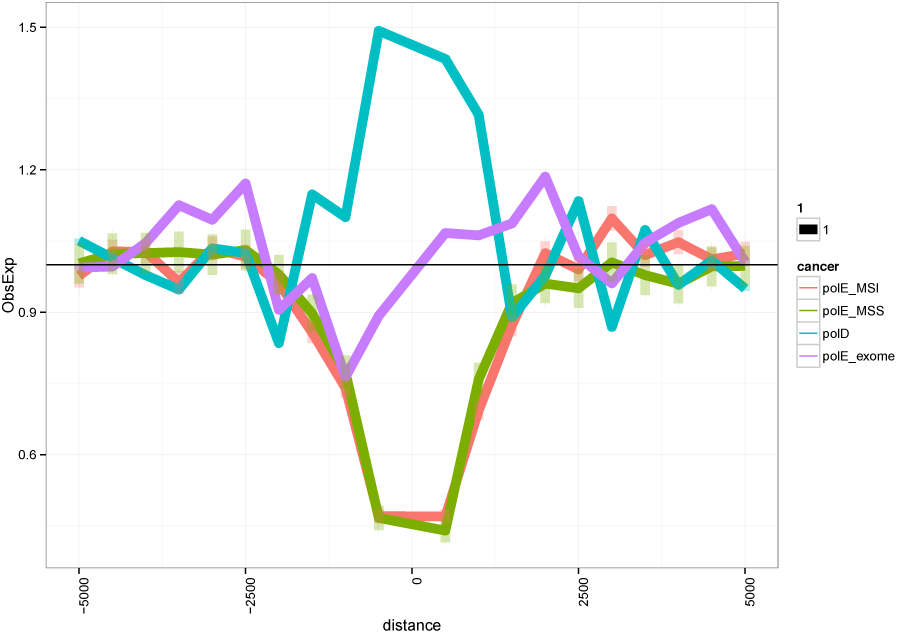
Somatic mutation rates on a 5-kilobase scale centered around potential human replication origins defined as ORC binding sites: in POLE* cancers with MSI (red line) and MSS status (green line); in POLE* tumours for which only exome data were available (blue line). To control that a mutation rate peak at replication origins in POLD* tumours couldn’t be explained by using only exome data, we provide a similar plot for POLE* tumours with only whole-exome sequences available (magenta line). No similar peak was observed.

**Figure S3.**
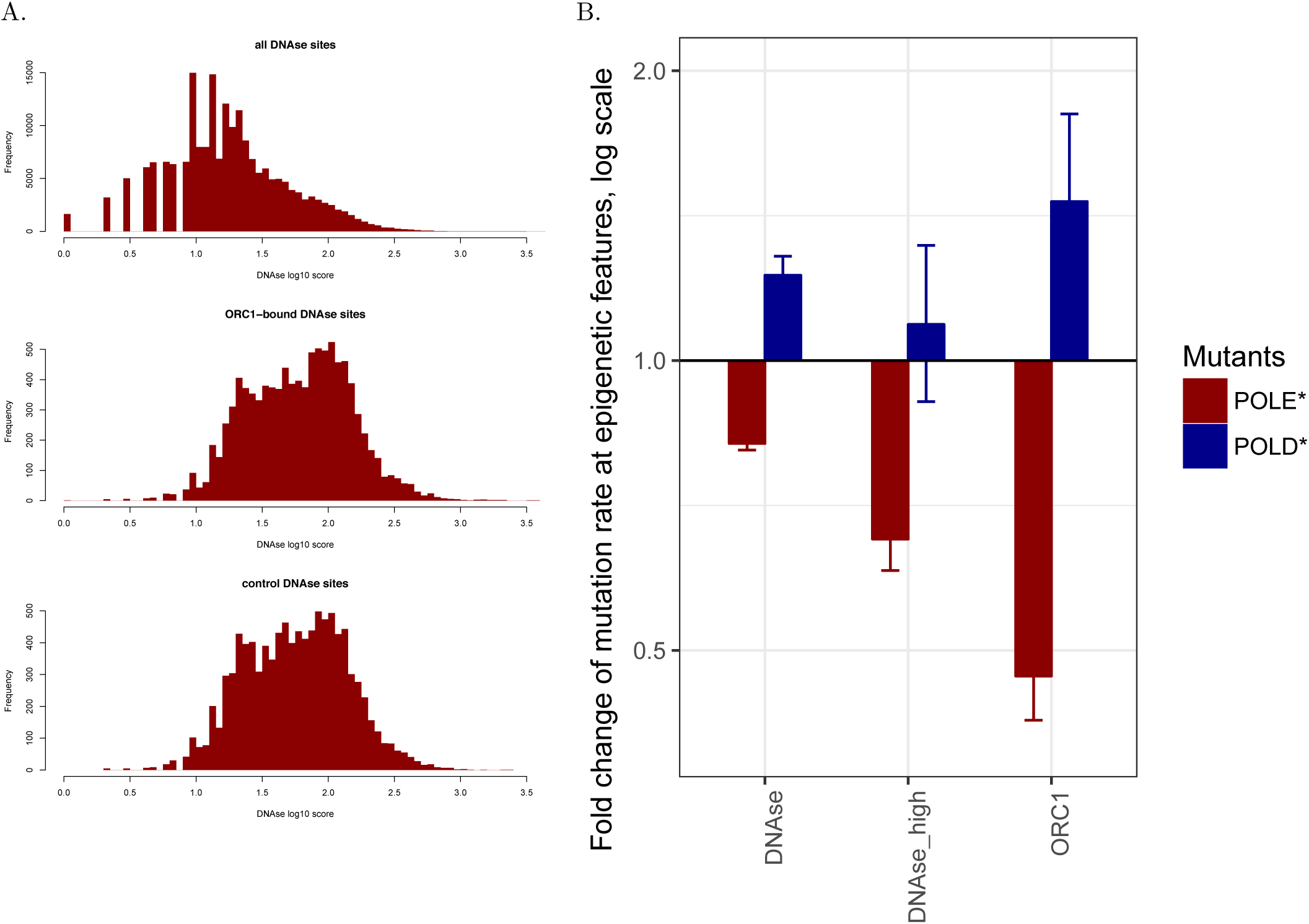
(A) Histograms showing the distributions of DNAse hypersensitivity scores in all DNAse hypersensitivity sites (DHS, top panel), in the DHSs overlapping with an ORC site (middle panel), in the control set of DHSs which did not overlap with ORC sites but had a similar distribution of DNAse hypersensitivity scores. (B) Similar to Figure 1A. For each epigenetic feature (x-axis), we plotted context-corrected mutation rate adjacent to this feature (-500..+500bp window) normalized by mutation rate further from the regions of interest (-5000..-4500 bp and 4500..5000 regions, see Methods section). DNAse high represents a subset of DNAse sites having the score distribution similar to that of the DNAse sites overlapping with ORC sites.

**Figure S4.**
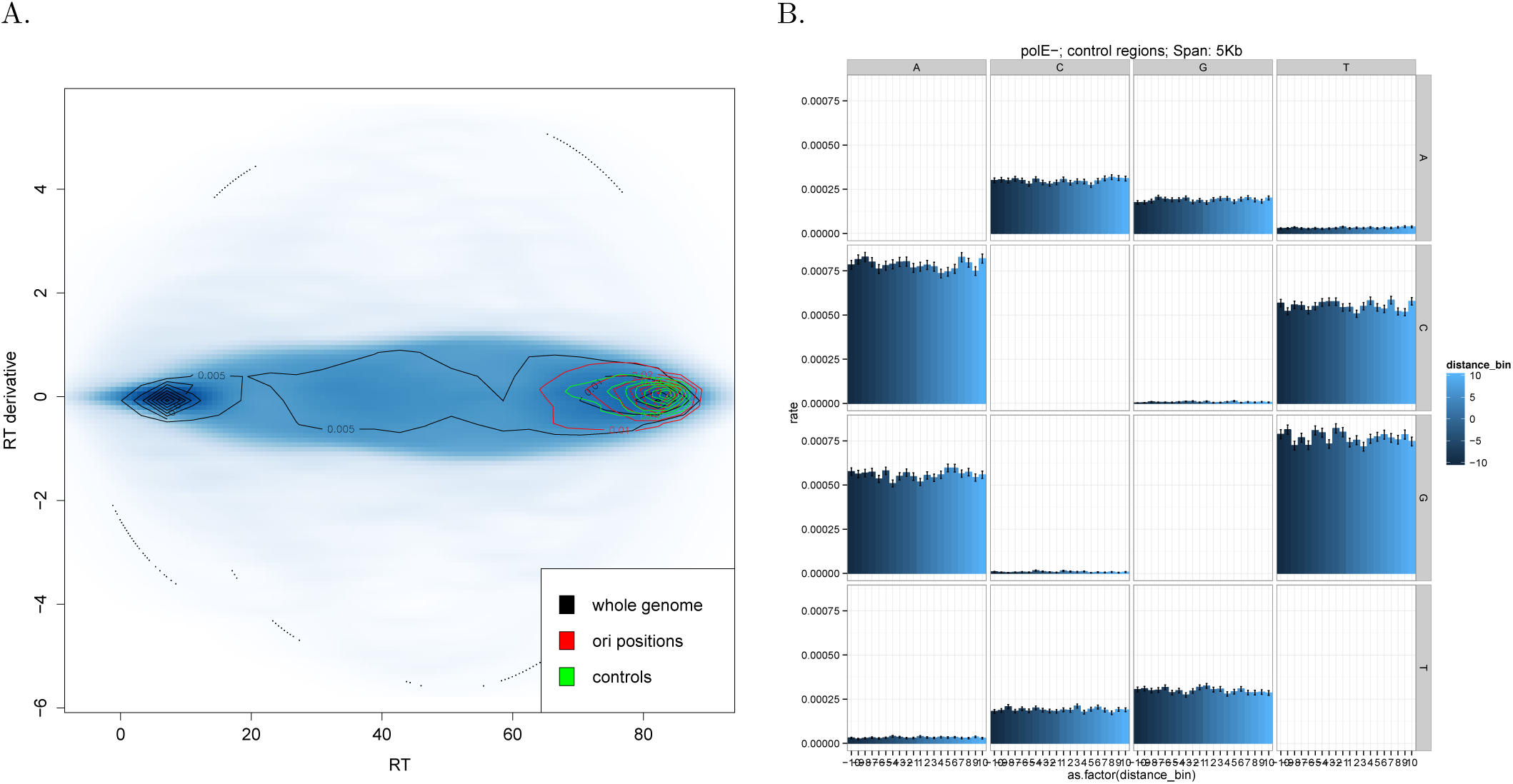
(A) Distribution of origins with respect to replication timing and derivative or replication timing. Replication timing derivative was calculated as a difference of replication timing values between consecutive kilobase bins. 2D distributions were depicted by contour plot. (B) Mutation rates on a kilobase scale in POLE* tumours centered around control set of genomic loci with replication timing similar to real origins.

**Figure S5.**
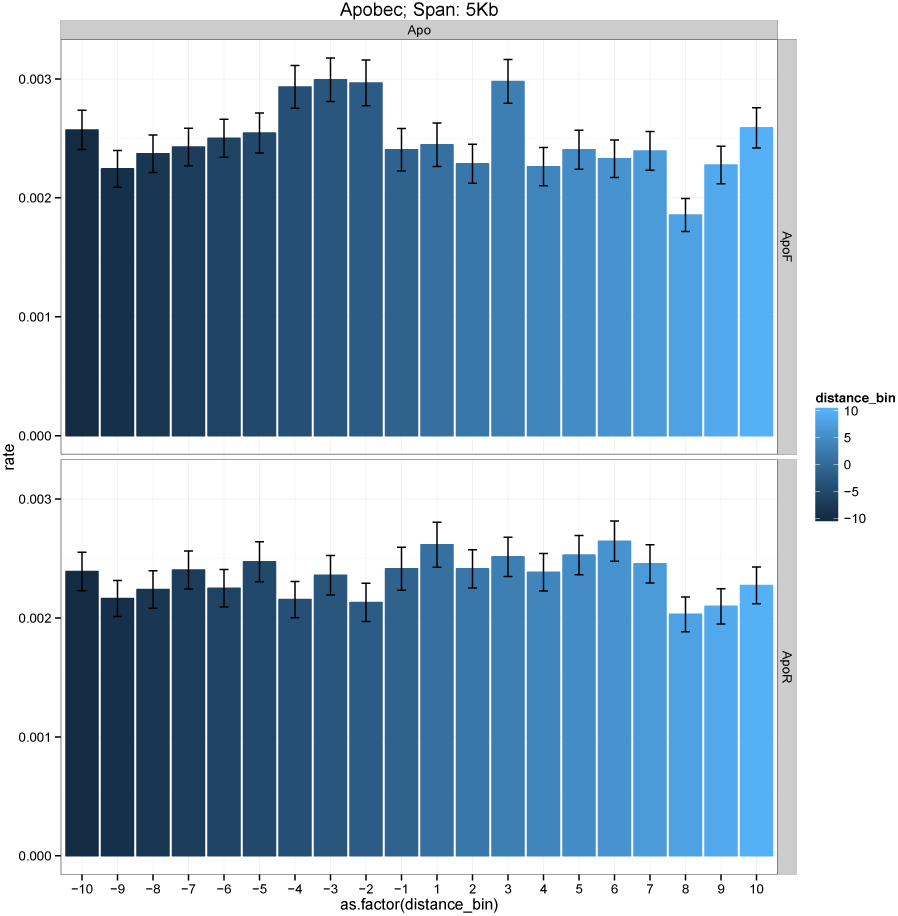
Somatic mutation rates on a 5-kilobase scale centered around potential human replication origins defined as ORC binding sites in tumours enriched with APOBEC-induced somatic mutations. The rate of APOBEC-induced mutation on forward and reverse strands was calculated as described in (Methods).

**Figure S6.**
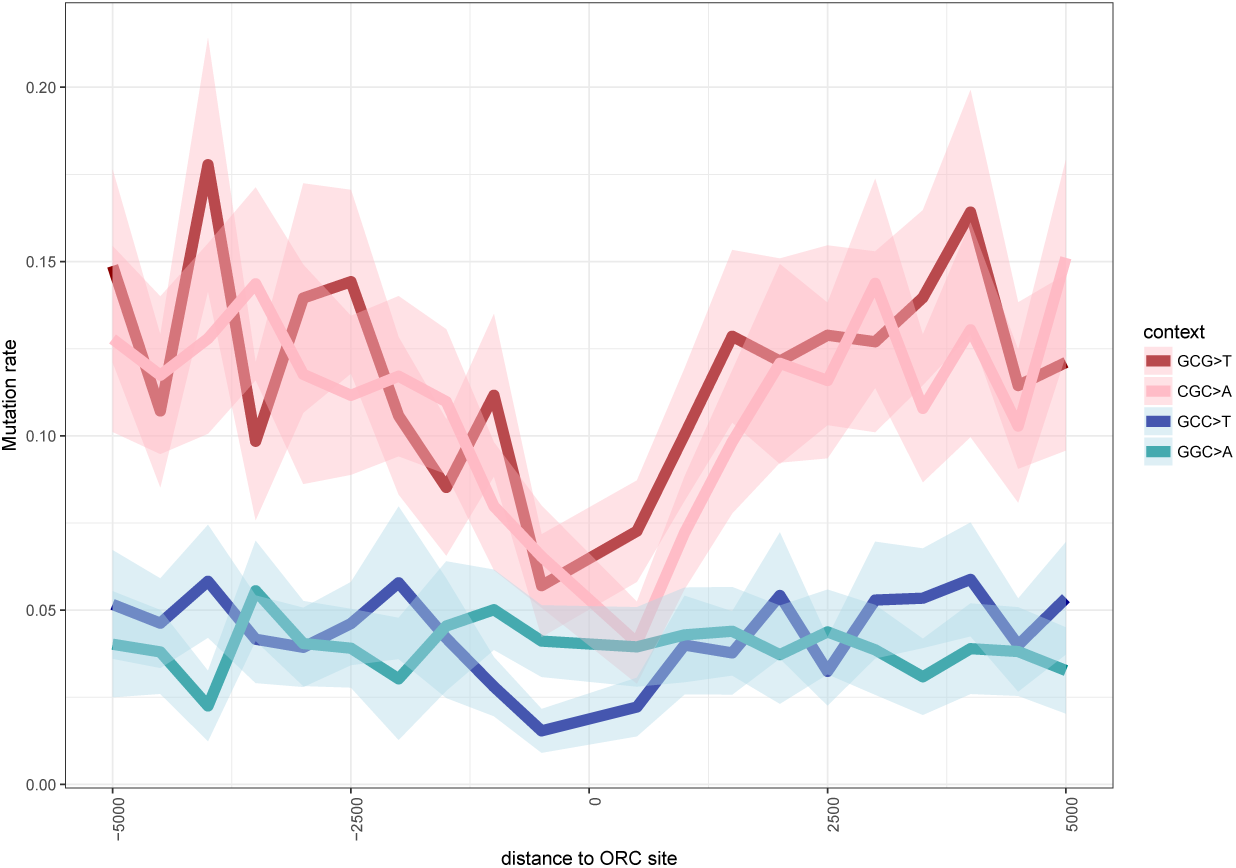
Context-corrected somatic mutation rates (see Methods) on a 5-kilobase scale centered around potential human replication origins defined as ORC binding sites in *POLE*^*wt*^/*POLD*^*wt*^ tumours in the contexts characteristic to polE* (GCG→T and its complement CG→A) and to polD* (GCC→T and its complement GGC→A).

**Figure S7.**
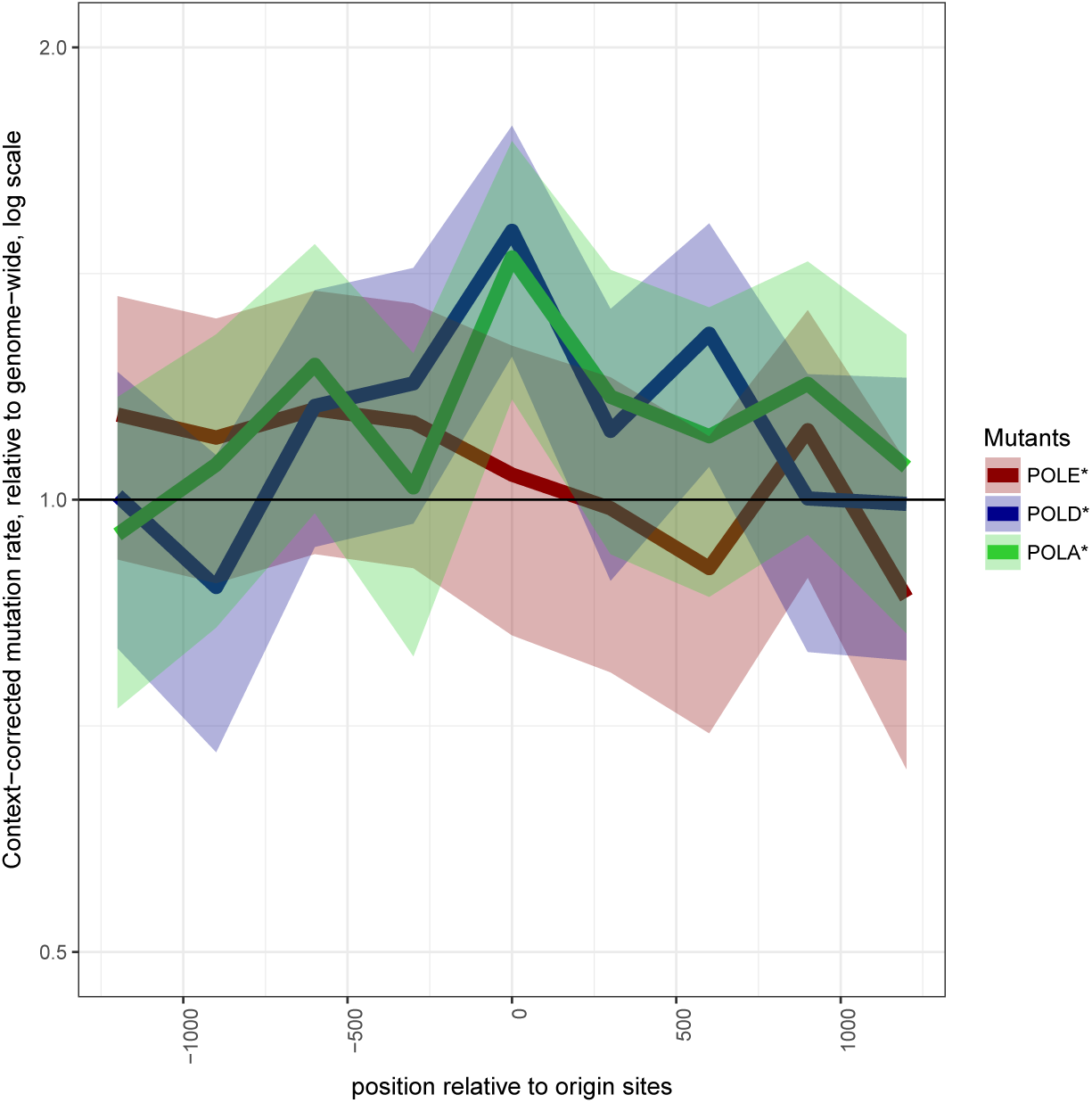
Similar to Figure 1b. Context-corrected somatic mutation rates (see Methods) centered around yeast replication origins in *pol2-M644G* (POLE*), *pol3-L612M* (POLD*) and *pol1-L868M* (POLA*) *Saccharomyces cerevisiae* strains.

**Figure S8.**
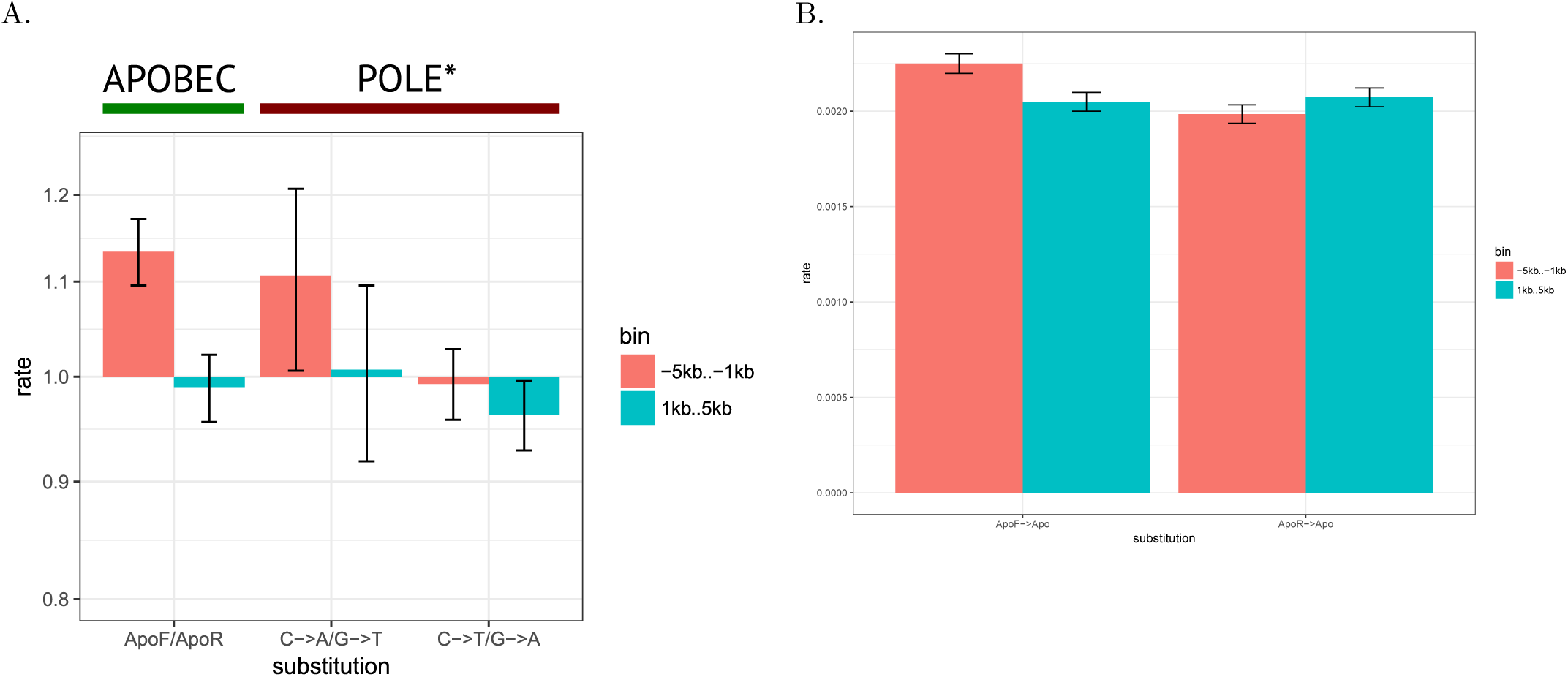
Mutational asymmetry of APOBEC-and POLE*-induced mutations at origins. (A) Ratios of mutation rates for complementary pairs of nucleotide substitutions (e.g., C→ A/G→ T) in cancer genomes enriched with APOBEC mutations and in POLE* MSI tumours to the left and to the right from an ORC site. Only regions further than 1kb but closer than 5kb from ORC sites were considered. Error bars indicate standard deviation. (B) The rate of APOBEC-induced somatic mutations on the forward and the reverse strands (see Methods) further than 1kb but closer than 5kb from ORC sites was counted to the left and to the right from an origin. Error bars indicate standard deviation.

**Figure S9.**
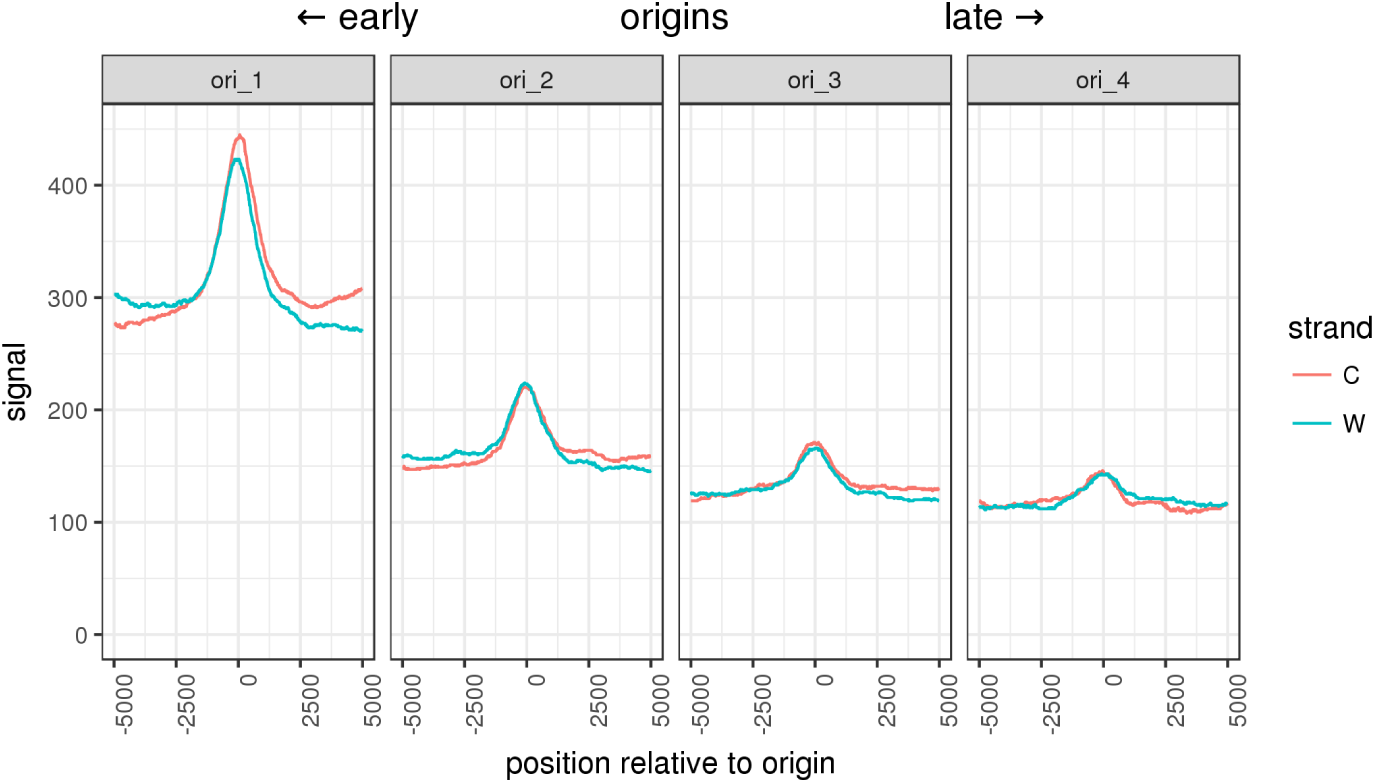
Density of Okazaki fragments profiled by OK-seq around ORC sites split into four groups according to their replication timing. The fragments mapped to Watson (W) and Crick (C) strands are plotted separately. The enrichments of short *de novo* synthesized DNA fragments at ORC sites exceeds the difference of Okazaki fragment density between the leading and the lagging replication strands.

**Figure S10.**
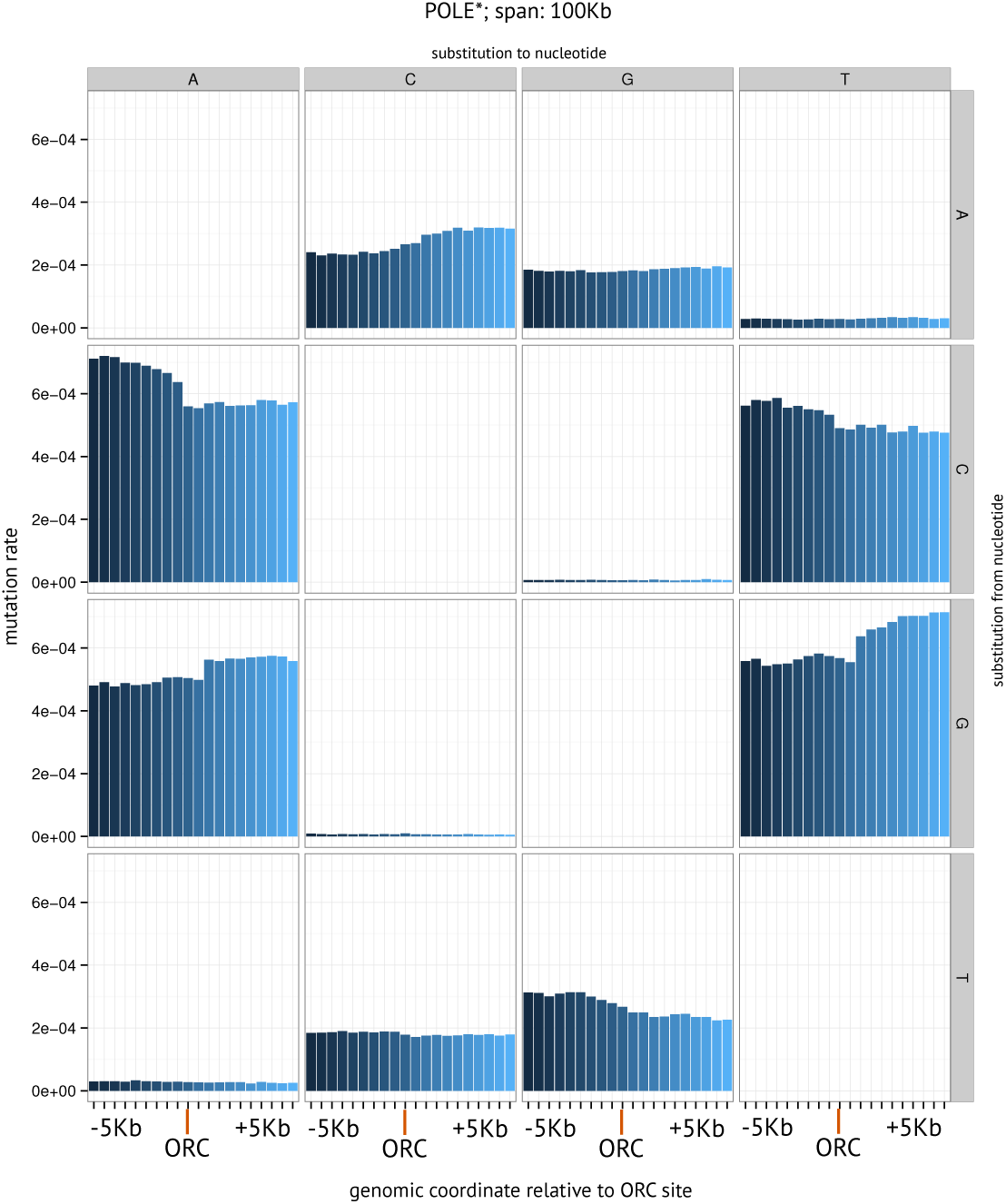
Mutation rates on a 100-kilobase scale centered around human replication origins in POLE* tumours. Notice an asymmetric pattern for the most of the substitutions.

